# A qPCR method facilitates study of absolute abundance, ecology, and inoculation fate of ciliate predators on the leaf surface

**DOI:** 10.64898/2026.08.14.744910

**Authors:** Stephen J. Taerum, Ravikumar R. Patel, Blaire Steven, Lindsay R. Triplett

## Abstract

Predatory protists are important in shaping terrestrial microbial ecosystems, but their roles in the phyllosphere, or the communities on aerial plant surfaces, are poorly understood. Previous work found that the order Colpodida dominated heterotrophic protist communities in the phyllosphere. While most protists were sporadically present, a few Colpodida variants were prevalent and abundant, indicating that these variants may represent species adapted to the phyllosphere. To identify these organisms, we cultured colpodids from field-collected tomato leaves and performed phylogenetic analysis of the 18S rRNA gene. Five of nine independent isolates matched the most prevalent Colpodida variant previously identified as leaf-enriched through amplicon sequencing, and these isolates comprised a novel clade of *Paracolpoda steinii.* When compared to a maize root isolate of *Colpoda inflata,* an abundant rhizosphere ciliate, a *P. steinii* isolate was similar in size and growth yield on *E. coli,* but grew to higher yields and formed large cyst clusters when incubated with model phyllosphere bacteria prey *Erwinia* and *Pseudomonas*. We developed and validated quantitative PCR (qPCR) methods for detection and cell abundance estimation of the *P. steinii* phyllosphere clade, *C. inflata*, and the order Colpodida in environmental samples. In inoculated greenhouse plants, qPCR-estimated protist populations matched measured inoculum levels, and protist inoculum was still detectable after five days. In an uninoculated tomato field, *P. steinii* was detected on all plants, with greatest abundances observed in lower leaves and after a rain event. *P. steinii* comprised up to 18.7% of total leaf Colpodida populations, which were estimated at up to ∼1400 organisms per gram of fresh weight. The findings demonstrate that Colpodida communities are consistently present on tomato leaves, dynamically affected by the abiotic environment, and include significant populations of *P. steinii*. We propose that the *P. steinii* isolates and qPCR tools presented can be used as a model system to investigate colonization and distribution patterns, biotic interactions, genetic adaptations, and agricultural applications of leaf predation.

## INTRODUCTION

Protists are an abundant and diverse component of the plant microbiome. There are an estimated 10^4^ - 10^8^ protists per gram of soil (Adl and Coleman 2005; Leach et al. 2017; Oliverio et al. 2020) representing hundreds to thousands of unique variants (i.e., operational taxonomic units or amplicon sequence variants) from all the major protist supergroups (Ceja-Navarro et al. 2021; Taerum et al. 2022; Zhang et al. 2021). Rhizosphere protists exert well-established roles in plant growth through cycling nutrients (Bonkowski 2004), shaping bacterial communities and disease outcomes (Asiloglu et al. 2020; Guo et al. 2022), cross-kingdom communication and gene expression (Jousset et al. 2010; Patel et al. 2025), and introducing bacteria to the rhizosphere (Taerum et al. 2024, 2025). Far less is known about the abundance, diversity and function of predatory protists in the phyllosphere, or aboveground surfaces of the plant (Leach et al. 2017). Following a similar pattern to phyllosphere bacterial communities, leaf protist communities are highly distinct from those in the rhizosphere, exhibiting reduced diversity and greater variability and stochasticity, with the exception of a few consistently observed core taxa (He et al. 2024; Taerum et al. 2023). There is evidence these simple protist communities have important ecosystem functions; they act as a reservoir for bacteria in the phyllosphere (Lin et al. 2023), shape microbial communities on leaves through selective predation (Flues et al. 2017), and form positive network associations with bacteria (Taerum et al. 2023). Despite this potential importance, a lack of information about the species identities, life cycle, and ecological adaptations of phyllosphere protists has prevented a detailed understanding of their roles.

Microscopic observations and relative abundance data from amplicon sequencing studies have suggested that protists in the order Colpodida (class Colpodea, supergroup Alveolata) dominate the phyllosphere. Colpodida is an order of cosmopolitan ciliates widely distributed in global soils, detritus, oceans, freshwater, and air (Canals et al. 2020; De Groot et al. 2021; Li et al. 2024; Oliverio et al. 2020). Soil Colpodida significantly improved the growth of plants when added to plant roots (Asiloglu et al. 2020; Zhang et al. 2022), introduced bacteria to the rhizosphere (Taerum et al. 2025) and facilitated bacterial transportation along plant roots (Hawxhurst et al. 2023; Micciulla et al. 2024). Early microscopic enumeration studies reported that aerial tissues of diverse plants were dominated by a species morphologically identified as *Colpoda cucullus* (Mueller and Mueller 1970). This protist was estimated to reach up to 800 organisms per gram of leaf tissue, and was described as an environmentally resilient protist that rapidly replicates in dew and then forms resting cysts during wet-dry cycles, traits well-suited to survival of the fluctuating phyllosphere environment (Bamforth 1973). Culture-independent studies later validated *Colpoda* and related ciliates as common leaf surface residents, although protists from other lineages including Amoebozoa and Rhizaria are also present on leaves (He et al. 2024; Sapp et al. 2018).

In a recent amplicon sequencing survey, we found that Colpodida dominated the heterotrophic protist communities of solanaceous leaves, with relatively few variants comprising over 50% of the reads (Taerum et al. 2023). We also observed signs that certain Colpodida might be phyllosphere specialists, as over a dozen sequence variants were exclusively detected in leaf but not root wash samples. Many of these variants were rare, but one Colpodida variant was present in high relative abundance in the majority of leaf samples while rarely detected in the rhizosphere. Based on this observation and on previous observations of *C. cucullus* dominance, we hypothesized that a small number of Colpodida species were widespread leaf residents with potential to impact phyllosphere ecology and plant health.

In this study, we sought to develop isolates and quantitative tools that would allow identification, enumeration and lab study of phyllosphere resident Colpodida. An isolation survey in tomato predominantly yielded isolates matching the prevalent leaf Colpodida variant, and these were placed in a novel clade in the species *Paracolpoda steinii*. A representative *P. steinii* isolate was characterized and grew to high yields when supplied with phyllosphere bacteria. We designed quantitative PCR (qPCR) protocols to quantify cells in the phyllosphere clade of *P. steinii* and across the order Colpodida, and found that the tests accurately estimated known inoculum populations on greenhouse-inoculated tomato leaves. The tests were used to enumerate Colpodida on leaves of tomato grown in a Connecticut field, finding that *P. steinii* and broader Colpodida communities were ubiquitous on tested plants, and populations varied dynamically by leaf age and wetness condition. These studies validate Colpodida as a leaf inhabitant of the tomato plant, and provide baseline observations and experimental resources for studying Colpodida populations in the phyllosphere.

## METHODS

### Isolation of Colpodida from tomato leaves

Leaf fragments were collected from flowering tomato plants (Mountain Fresh) grown at Lockwood Farm in Hamden, CT, USA, on August 1, 2022. The fragments were placed in sterile Petri dishes containing 5 mL autoclaved dH_2_O. The plates were sealed with parafilm and stored in the dark for 48 hours. The plates were examined for activity at 100× magnification every 24 hours using a Zeiss ID02 Invertoscope inverted microscope. Active cells with characteristic Colpodid morphology were individually collected from the solution using glass capillaries attached to a suction bulb. Single cells were transferred to 1 mL PAGE’s solution (0.12 g of NaCl, 0.004 g of MgSO_4_•H_2_O, 0.004 g of CaCl_2_, 0.142 g of Na_2_HPO_4_, and 0.136 g KH_2_PO_4_ in 1 L dH_2_O), amended with heat-killed *E. coli* DH5α (OD_595_ = 0.005) and grown in 24-well plates. When replication of a single morphotype was observed in a well, 1 mL of culture was added to a 50 mL polypropylene tube containing 9 mL of PAGE’s solution amended with heat-killed *E. coli*. The cultures were subcultured every 6 weeks by transferring 1 mL of culture to 9 mL of sterile PAGE’s solution containing heat-killed *E. coli*.

### 18S rRNA gene sequencing of Colpodida

Genomic DNA was extracted from 250 µL of active protist culture using the ZymoBIOMICS™ DNA Miniprep Kit (Zymo Research, Tustin, CA, USA). Nearly complete 18S rRNA genes (∼1,781 bp) were amplified using the primers EukA (5’-AACCTGGTTGATCCTGCCAGT -3’) and EukB (5’-GTAGGTGAACCTGCAGAAGGATCA-3’; Medlin et al. 1988). PCR reactions were comprised of 1× DreamTaq DNA Polymerase (ThermoFisher Scientific, Waltham, MA, U.S.A.), 0.5 mM of each primer, and 2 ng of template in 25 µL aqueous reactions. PCR conditions were as follows: 3 min at 94°C, followed by 30 cycles of 94°C for 45 s, 60°C for 30 s, and 72°C for 90 s, followed by a final extension of 72°C for 10 min. Amplification products were cloned into the One Shot™ TOP10 vector per manufacturer’s instructions (ThermoFisher Scientific) and sequenced by Sanger sequencing using the M13F (5’-GTAAAACGACGGCCAG -3’) and M13R (5’-CAGGAAACAGCTATGAC -3’) primers at the Keck DNA Sequencing Core (Yale University, New Haven, CT, USA).

### Phylogenetic analysis

Sequences of the nine phyllosphere protists obtained in this study were aligned with 52 sequences obtained from Genbank: ten sequences of rhizosphere Colpodida from Taerum et al. (2022), 37 from Colpodida that were listed in the curated Protist Ribosomal Reference (PR^2^) collection, and five from the order Cyrtolophosidida as an outgroup (Table S1). Sequences were aligned using the Auto strategy in MAFFT v. 7 (Katoh and Standley 2013), and then manually checked for regions of ambiguous alignment. The alignment was trimmed to 1714 bp. Mean uncorrelated pairwise distances (p-distances) between the sequences were calculated using MEGA v. 7.0.26 (Kumar et al. 2016).

Maximum likelihood (ML) and Bayesian inference (BI) analyses were used to determine the phylogenetic positions of the Colpodida isolates. ML analyses were conducted using raxML v. 8.2.10 (Stamatakis 2014), following a GTR+Γ evolutionary model. Bootstrap support was obtained using 1000 ML replicates. BI analyses were conducted using Mr. Bayes v. 3.2.7 (Ronquist et al. 2012), following a GTR+Γ evolutionary model. Four Markov chain Monte Carlo chains were run with 5,000,000 generations, and posterior probabilities were calculated by sampling every 100 generations. Tracer v. 1.7.2 (Rambaut et al. 2018) was used to calculate burn-in, and trees generated in the burn-in phase were discarded.

### Comparison of protist sequences with amplicon sequences from previous study

The V9 regions of the 18S rRNA genes of 11 Colpodida that were enriched or de-enriched in the phyllosphere of solanaceous plants (Taerum et al. 2023) were downloaded from the National Center for Biotechnology Information (NCBI; BioProject PRJNA816421). These sequences were aligned with the phyllosphere sequences obtained in the present study using MAFFT. The sequences were then visually compared in MEGA to identify any matching sequences.

### Characterization of protist isolates

Active and encysted cells from 48-hour old cultures were photographed and measured using a Zeiss Axiovert A1 inverted microscope and an Axiocam 202 mono camera at 1000× magnification under oil immersion and brightfield imaging. The population growth and encystment behaviors of the two strains were then compared. To wash residual bacteria, polypropylene tubes containing 20 mL of encysted protists were centrifuged at 1,500 g for 20 minutes. Approximately 15-17 mL of supernatant was removed, after which the protist cells were washed with 15 mL of fresh PAGE’s solution. This process was repeated twice, after which the cells were diluted to ∼1000 cells per mL in 24-well plates with wells containing 1 mL of PAGE’s solution amended with heat-killed *E. coli* (OD_595_ = 0.05). Five wells were set up for each culture, and the plates were incubated for 1 week at 22°C in the dark. Population sizes were estimated by manually counting active and encysted cells from three photos taken in a triangular pattern with an Axiocam 202 mono camera at 200× magnification every 24 hours.

### Protist co-culture with plant pathogenic bacteria

Protist cultures of *P. steinii* isolate CLRT4A and *Colpoda inflata* maize isolate UC22 were prepared by 1/10 dilution of encysted cultures into PAGE’s solution amended with heat-killed *E. coli* (OD_595_ = 0.01) and room temperature incubation for three days. Cultures were diluted with PAGE’s solution to a density of 400 protist cells per mL. Bacterial cultures (*Pantoea agglomerans* 299R (Remus-Emsermann et al. 2013), *Pseudomonas savastanoi* pv. *phaseolicola* 1448A (Arnold et al. 2011), *Erwinia amylovora* Ea88 (McManus and Jones 1994), *Dickeya dadantii* 3937 (Glasner et al. 2011), *Dickeya chrysanthemi* AC4150 (Chatterjee et al. 1983), *Dickeya zeae* QZ933) were grown from single colonies by shaking incubation overnight at 28°C in M9 minimal medium (Maniatis et al. 1982) amended with 0.4% glucose and 1 mM thiamine HCl, then washed and resuspended to an OD_600_ of 0.2 in the same medium. Heat-killed *E. coli* was resuspended to the same optical density as an inert prey control. 500 µL protist culture and 500 µL bacterial culture were combined in the wells of 24-well plates. Therefore, each well contained 100 protists and a bacterial OD_600_ of 0.1 in 1 mL of PAGE’s solution/M9 medium mixture. Plates were incubated in the dark at 22°C for 5 days, after which protists had stopped actively swimming. Protists were counted using an inverted microscope as described above. For wells containing large clusters of cysts (i.e. CLRT4A incubated with *E. amylovora* or *P. syringae* pv. *phaseolicola*), well contents were vortexed in an Eppendorf tube and diluted 1/10 before counting.

### qPCR primer design

Primers based on the 18S rRNA genes of *P. steinii* CLRT4A and *C. inflata* UC22 were designed using Primer-BLAST (Ye et al. 2012). Amplicon length range was filtered to 75-300 bp, and melting temperature range was set to 50°-65°C, with a maximum T_m_ difference of 5°C. The primers were checked against the nt database, and primers that matched non-target Colpodida were excluded. Targets with 1 or more mismatches were ignored. The top scoring 100 primer pairs from each set were screened using the PR^2^ primer database (Vaulot et al. 2022), and primers were excluded if they matched the sequence of a non-target protist, allowing for no mismatches. Primers were designed based on an alignment of *C. inflata* and *P. steinii* isolates, with no taxa excluded using Primer-BLAST. For Colpodida primers top scoring 200 primer pairs were screened using PR^2^ to maximize the number of Colpodida amplified and minimize the number of non-Colpodida, allowing for no mismatches.

Specificity of candidate primers were tested by amplification against DNA from the target isolates and other diverse protists in our collection (Taerum et al. 2022): *Flamella* (Amoebozoa; UC122), *Ochromonas* (Stramenopiles; UC48), *Cercomonas* (Rhizaria; UC19), *Thaumatomonas* (Stramenopiles; UC70), and tomato leaf. PCR reactions were conducted as described above.

### Standard curves to calculate gene copies per cell

The 18S rRNA gene of isolates UC22 and CLRT4A were amplified using the primers EukA and EukB, following the PCR protocol listed above. Amplification products were purified using the GeneJET Gel Extraction and DNA Cleanup Micro (ThermoFisher Scientific) kit and quantified using a Qubit Fluorometer (ThermoFisher Scientific). Based on the concentration values, DNA was serially diluted to concentrations of 10^1^ to 10^6^ 18S rRNA gene amplicons/µL. qPCR reactions were conducted on a CFX96 Touch Real-Time PCR machine (Bio-Rad, Hercules, CA, U.S.A.). Reactions contained 1× SsoAdvanced Universal SYBR Green Supermix (Bio-Rad), 0.5 mM forward and reverse primers, and 2 ng of template in a 10 µL volume. The cycling protocol started at 2 min at 95°C followed by 40 cycles of 95°C for 10 s and 60°C for 15 s. Reactions were conducted in triplicate. Standard curves were also prepared from cultures of UC22 and CLRT4A adjusted to contain 10^1^, 10^2^, 10^3^, and 10^4^ active cells in 250 µL of PAGE’s solution (five replicates each), to correlate direct cell counts with quantified gene copy number. The protocol was evaluated on encysted CLRT4A cells to determine its applicability across distinct physiological states.

### Tomato inoculation

Tomato seeds (Mountain Fresh variety, Johnny Selected Seeds) were germinated in PROMix potting soil in 30 mm diameter inserts in the greenhouse in April, 2024 (16 h light, and temperatures ranged from 20°C – 28°C). After 28 days they were transplanted to 105 mm diameter pots containing potting soil and allowed to grow for an additional 14 days. Plants were watered every 2-3 days with autoclaved deionized water. On May 24, 2024, the tops of the terminal leaflets of the two newest leaves were spot inoculated with 10 µL of PAGE’s solution containing 1000 active cells of *P. steinii* isolate CLRT4A or *C. inflata* isolate UC22, or with sterile PAGE’s solution as a control. Seven plants were inoculated per treatment. After 1 hour, one inoculated leaflet was collected from each plant. The other inoculated leaflet was collected five days later. Leaf samples were weighed and macerated by sterile mortar and pestle in liquid nitrogen, and total DNA was extracted using the ZymoBIOMICS™ DNA Miniprep Kit. qPCR reactions were conducted on the sample DNA and standard curve DNA as described above using the three primer pairs. Reactions with a threshold cycle greater than or equal to 30 were considered undetected, but set to a sample count of 1 to allow for log conversions. Cell numbers were estimated using back calculations from Fig. 3F.

Two-way analysis of variance (ANOVA) was used to examine the impacts of inoculum (CLRT4A, UC22, or PAGE’s solution) and time after inoculation (24 h vs. 5 days) on cell counts estimated with each primer pair. In addition, paired t-tests were used to compare cell enumeration between specific and broad primers. All statistical analyses were conducted using R v. 4.4.2.

### Field sampling

Tomato leaves of variety Sungold (Tokita) were sampled from Lockwood Farm in Hamden, CT during July 2023. Sampling began 87 days after seeding and 41 days after planting in the field, when plants were roughly 5 feet tall and in green fruit formation. Water was provided by drip irrigation. Dry leaf samples were collected July 13 at midday when the temperature was 32.3°C; the previous three days had reached 30°C or higher and had no cloud cover or precipitation. Moist leaf samples were taken on the morning of July 17^th^, at 27.2°C in misty and overcast conditions, 11 hours after the end of 14 hours of intermittent rain culminating in 7.7 cm of precipitation. For each timepoint, two leaves were collected from each of five plants, or two plants chosen at random from each of two rows, and one plant chosen from the third row. One of the top two fully expanded leaves were sampled and trimmed to 5 leaflets starting with the terminal leaflet. One of the bottom-most leaves was then sampled and trimmed to 3 leaflets including the terminal leaflet. Selected bottom leaves were neither senescing, touching the ground or other plants, or extending well into the row. Corresponding leaf samples were collected from the same five plants at both timepoints. Leaves were collected with flame-sterilized scissors and placed in pre-weighed polypropylene tubes, and placed immediately on ice. Within one hour of sampling, tubes containing leaves were weighed and stored at -80°C until DNA extraction. DNA extraction and qPCR amplification were performed following the same protocol as used in the inoculation experiment described above. Colpodida cell number estimates were back calculated from the curves in Fig. 3F.

Two-way ANOVA was used to examine the impacts of recent precipitation (wet vs. dry), leaf location (top vs. bottom), and the interaction between the two independent variables on cell counts estimated with each primer pair. Paired t-tests were used to compare cell counts between broad and specific primers. All statistical analyses were conducted using R.

## RESULTS

### Isolation and phylogenetic placement of Colpodida isolates from the tomato phyllosphere

We previously found that a small number of Colpodida variants dominated the distinct predatory protist communities on solanaceous leaf surfaces (Taerum et al. 2023). To enable better understanding of leaf Colpodida ecology, we aimed to isolate protists that were representative of leaf-enriched variants. Nine protists with distinctive Colpodida “kidney shape” morphology (Foissner et al. 2011) were independently isolated from the leaves of field-grown tomato by single-cell isolation from leaf washes, and were maintained in PAGE’s solution fed with heat-killed *Escherichia coli*. Five isolates from four plants shared identical 18S rRNA V9 region sequence with the most prevalent phyllosphere-enriched Colpodida amplicon sequence variant identified in our previous work (Taerum et al. 2023)(Fig. S1, Table S1). To identify the isolates, we compared near full-length 18S rRNA gene sequences with those of previously isolated Colpodida from the maize rhizosphere (Taerum et al. 2022) and other published sequences.

Phylogenetic and p-distance analyses placed six isolates in a novel well-supported clade within the species *Paracolpoda steinii* (green shaded box in Fig. 1, p-distances in Table S2). Two isolates, CLRT16E and CLRT17A, were placed in another clade of *P. steinii* shared with isolates from the maize rhizosphere (Taerum et al. 2022). Other maize root Colpodida isolates from Taerum et al. (2022) were most closely related to *Colpoda inflata* or *Colpoda cucullus* (Fig. 1). Based on these observations, we determined that the leaf-enriched Colpodida variant was likely represented by our isolates.

**Figure 1:**
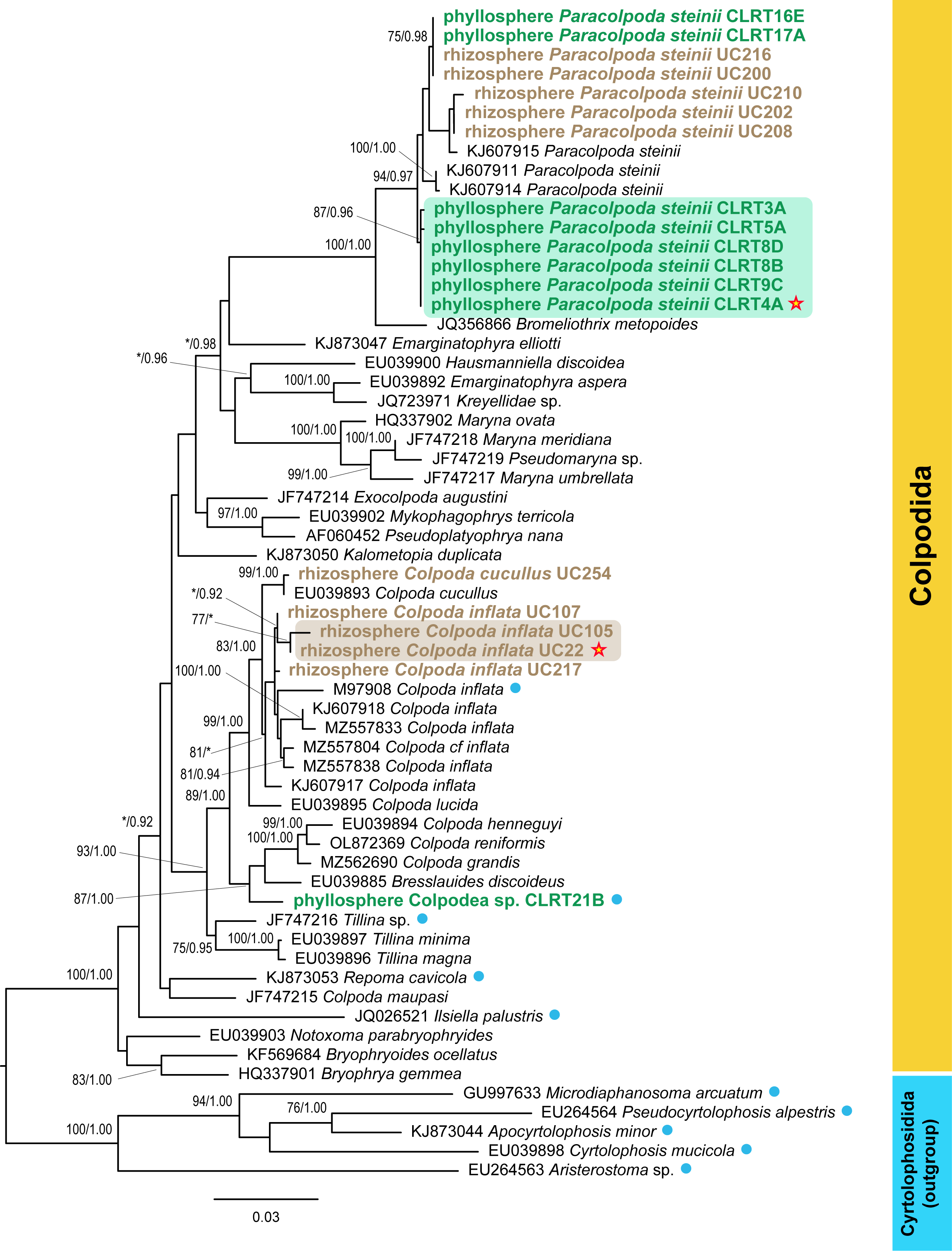
Maximum likelihood (ML) phylogram of Colpodida species based on most of the 18S rRNA gene. The tree contains sequences from nine phyllosphere-isolated Colpodida obtained in this study (colored green), ten rhizosphere-isolated Colpodida from Taerum et al. (2022; colored tan), 37 previously published sequences from Colpodida, and five sequences from Cyrtolophosidida as an outgroup. Each strain is indicated by its Genbank accession number or culture collection number (for the sequences obtained in this study) and its species name. ML bootstrap support based on 1000 pseudoreplicates and Bayesian inference posterior probabilities are shown at the nodes if greater than 75 and 0.9 respectively. The cultures used in this study for primer development and experimental inoculation are indicated with red stars. Taxa highlighted with green have target regions that 100% match the sequences of the primers 4A_2F/4A_2R, while taxa highlighted with tan have target regions that 100% match the sequences of the primers UC22_1F/UC22_1R. Blue circles indicate taxa that do not 100% match the target regions of the broad primers All_colp_4F/All_colp_4R.

### Characterization of growth and feeding differences between Paracolpoda steinii isolate CLRT4A and rhizosphere Colpoda inflata isolate UC22

We examined morphological and life cycle traits of *P. steinii* leaf isolate CLRT4A and C. inflata root isolate UC22 when grown in PAGE’s solution with heat-killed *E. coli*. *Paracolpoda steinii* and *C. inflata* were similar in appearance based on brightfield pictures, except that active *P. steinii* cells were longer and wider than active *C. inflata* (Table 1). CLRT4A and UC22 showed similar life cycles, with cysts becoming active within 24 hours of media inoculation and reaching peak activity at day 3 (Fig. S2, Table S3). However, UC22 reached higher density in culture under these conditions (Table S3).

**Table 1:**
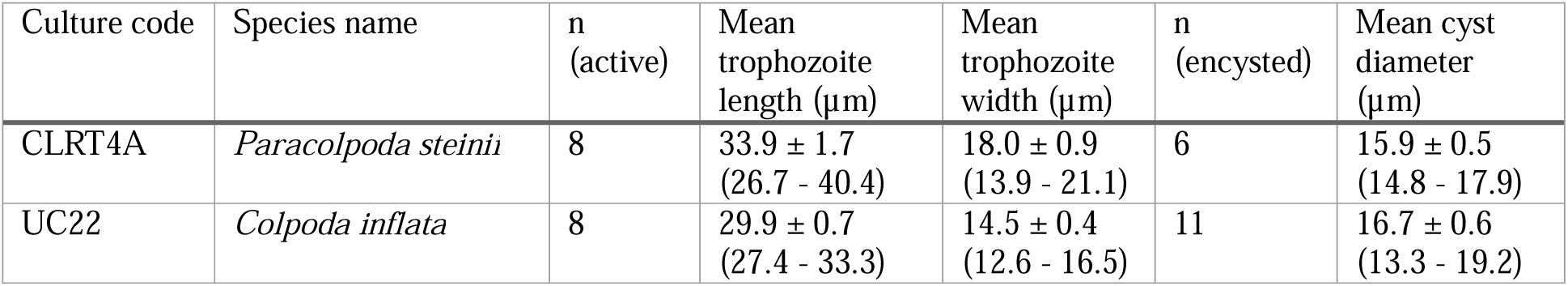
Mean cell sizes and standard errors of active and encysted P. steinii (CLRT4A) and C. inflata (UC22). Ranges are shown in parentheses after the means.

Given the leaf and root-specific enrichment of the *P. steinii* and *C. inflata* clades (Taerum et al. 2023), we asked whether isolates would exhibit different feeding patterns on plant pathogenic bacteria. Protist yield in the presence of a bacterial strain is an indicator of the bacteria’s suitability as a food source, while suppression of protist growth can indicate bacterial defense (Amacker et al. 2020); we hypothesized that *P. steinii* would be able to grow on phyllosphere bacteria. Isolates CLRT4A and UC22 were incubated with foliar pathogenic bacteria known to have epiphytic life stages (*E. amylovora*, *P.* savastanoi pv. *phaseolicola*; Lindow and Brandl 2003), root pathogenic species capable of soil survival (*Dickeya* spp.; Zhou et al. 2024), and a species that grows in either environment (*P. agglomerans*), and enumerated protist cells after five days. Although UC22 grew to greater density than CLRT4A on heat-killed *E. coli*, CLRT4A grew to 88% and 220% higher median density relative to UC22 when incubated with *E. amylovora* or *P. syringae* pv. *phaseolicola*, respectively (Fig. 2A). CLRT4A also showed a highly distinctive encystment phenotype in the presence of *E. amylovora* and *P. syringae*, forming large clusters of cysts not seen in UC22 cultures (Fig. 2B). In contrast, CLRT4A grew to lower density than UC22 in *D. zeae*, and the two protist isolates did not differ significantly in growth on *D. chrysanthemi*. *D. dadantii* and *P. agglomerans* supported little growth of both protists, suggesting that these pathogens have broad-spectrum predation defense (Fig. 2A). Dilution plating confirmed that addition of CLRT4A reduced bacterial density in wells of *E. amylovora*, but not in wells of *D. dadantii* or *P. agglomerans,* indicating that protist growth corresponded to a reduction in the pathogen population (Fig. S3). These results demonstrate that Colpodida isolated from different plant niches differ in their prey interactions among plant pathogens. Leaf-isolated *P. steinii* thrived to the greatest extent in the presence of two foliar pathogens, reaching high densities and forming large cyst clusters.

**Figure 2:**
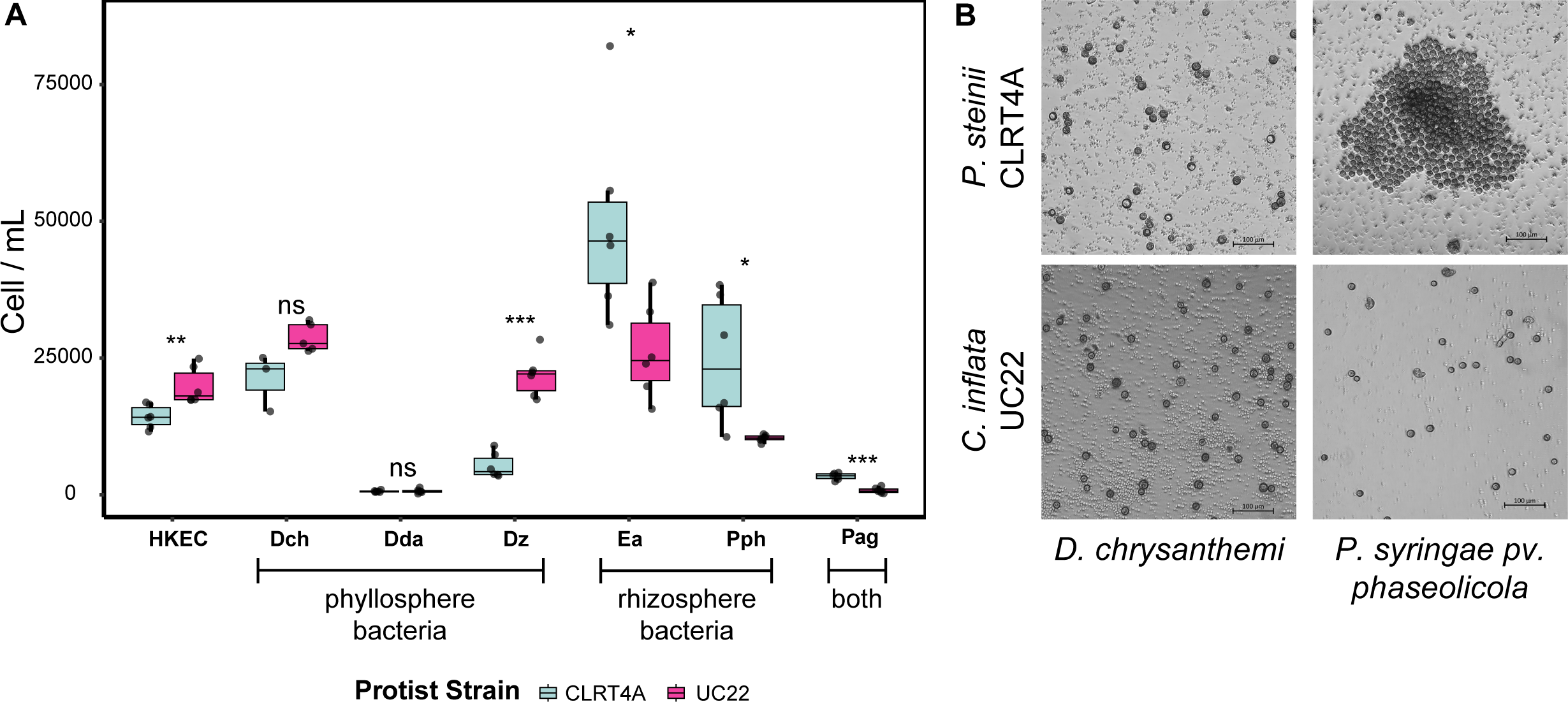
**A**. Growth of *P. steinii* isolate CLRT4 and *C. inflata* isolate UC22 in five sdays of co-culture with plant pathogenic bacteria. Asterisks indicate differences between protist isolates in an unpaired t-test (* p < 0.05 to *** p < 0.0005, n = 6). HKEC – heat-killed *E. coli*, Dch – *Dickeya chryanthemi*, Dda – *Dickeya dadantii*, Dz – *Dickeya zeae*, Ea – *Erwinia anylovora*, Pph – *Pseudomonas syringae* pv. *phaseolicola*, Pag – *Pantoea agglomerans*. **B.** Typical encystment pattern observed on incubation of CLRT4A (top) or UC22 (bottom) with *D. chrysanthemi* (left) and *P. syringae* pv. *phaseolicola* (right).

### Design of qPCR methods for environmental detection and quantification of leaf-enriched *P. steinii* and other Colpodida

While Colpodida have long been proposed as abundant residents of plant surfaces, absolute quantification methods are needed to understand the biotic and biotic drivers of their abundance. We designed qPCR primer pair 4A_2F/2R to target the phyllosphere-associated clade of *P. steinii* (Fig. 3A-B, green box in Fig. 1). To enable qPCR studies in the context of other Colpodida, we also designed two other primer sets: the first matches 91% of described Colpodida species in the PR^2^ database (Colp_4F/4R), and the second set amplifies *C. inflata* isolate UC22 (UC22_1F/1R; Fig. 3C-D). Isolate UC22 abundantly colonizes the maize rhizosphere and is an emerging model for bacterial-predator interactions in our studies (Patel et al. 2025; Taerum et al. 2024, 2025). Primers were validated to amplify DNA from the target organisms, but not to non-target Colpodida isolates or other protists (Fig. S4). We found a strong linear relationship between template DNA concentration and cycle threshold for all primer pairs (r^2^ ≥ 0.981, Fig. 3E). We next measured the number of 18S rRNA copies in different concentrations of active and encysted Colpodida. With active cells, we observed a strong linear relationship between cell number and target gene copy number between 10 and 10,000 cells per replicate (r^2^ ≥ 0.925, Fig. 3F). However, for cysts of *P. steinii* isolate CLRT4A, the number of 18S copies detected dropped dramatically relative to active cells (by 56 to 19,000-fold). The detected copy numbers increased in a linear fashion from 10^1^ to 10^3^ cells, but dropped sharply at 10^4^ cells (r^2^ = 0.388, Fig. S5). In light of this observation, we determined that the qPCR tests are target-specific and quantitative in measuring active cells, but likely do not have quantitative capacity to measure dormant Colpodida populations. Environmental samples are likely to contain Colpodida at different life stages, but Colpodida decrease in 18S copy numbers from peak activity to resting cyst stages (Zou et al. 2021). Henceforth we interpret quantitative cell measurements as equivalent measurements of the active population, or active cell equivalents (ACE), even if not explicitly stated.

### Detection of Colpodida following leaf inoculation

The qPCR tests were evaluated on tomato leaves after inoculation with Colpodida isolates in a greenhouse setting. Seven replicate leaves were inoculated with approximately 1000 active cells of *P. steinii* isolate CLRT4A or *C. inflata* isolate UC22, as measured by microscopic enumeration, and populations were estimated by the qPCR tests at 1 hour and 5 days. At 1 hour, qPCR tests estimated a mean ACE of 946 (s.e. = 137) *P. steinii* cells and 1209 (s.e. = 184) cells of all Colpodida on leaves inoculated with CLRT4A, closely matching the inoculum dose, while no *C. inflata* signal was detected (Fig. 4A; values in Table S4, S5). Similarly, a mean ACE of 746 (s.e. = 138) of *C. inflata* and 673 (s.e. = 117) of Colpodida were estimated on plants inoculated with *C. inflata* isolate UC22, but no *P. steinii* PCR signal was detected (Fig. 4B, Table S4).

**Figure 3:**
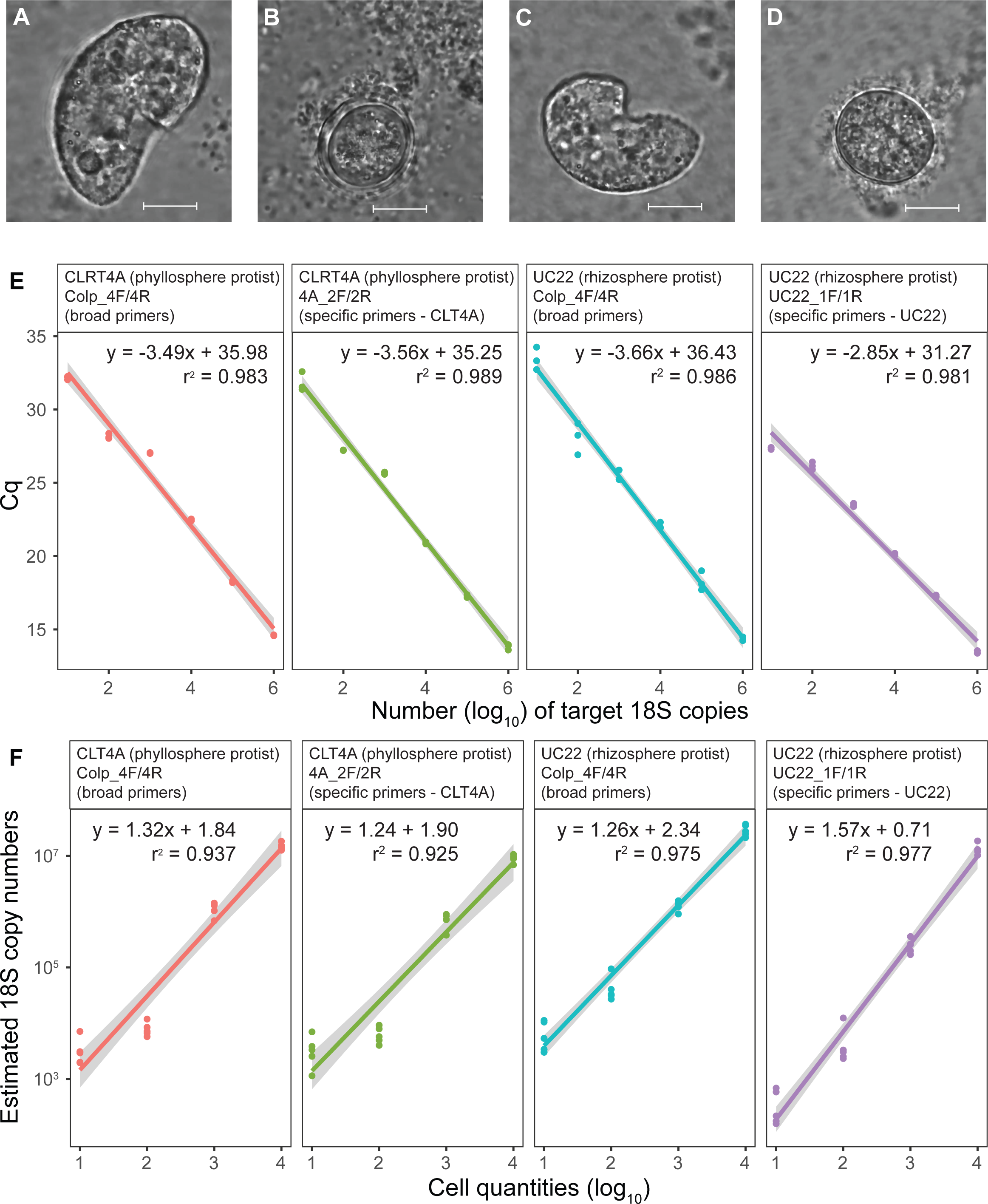
Standard curves generated during Colpodida qPCR primer development. **A-D**. Photos of the study organisms. Scale bars represent 10 µm. A and B are of active (A) and encysted (B) CLRT4A (*P, steinii*), while C and D are of active (C) and encysted (D) UC22 (*C. inflata*). **E**. Standard curves showing the relationship between the concentrations of 18S fragments and the threshold cycle at which amplicons were detected (n = 3). **F**. Standard curves showing the relationship between cell concentrations and estimated numbers of 18S fragments in the samples (n = 5). For E-F, the standard curves are shown for the specific primers on the target protist, and the broad primers on both protists. The lines represent the best fits based on the linear equations, which are shown above the lines along with the coefficients of determination. Grey shading represents the 95% confidence intervals around the line.

**Figure 4:**
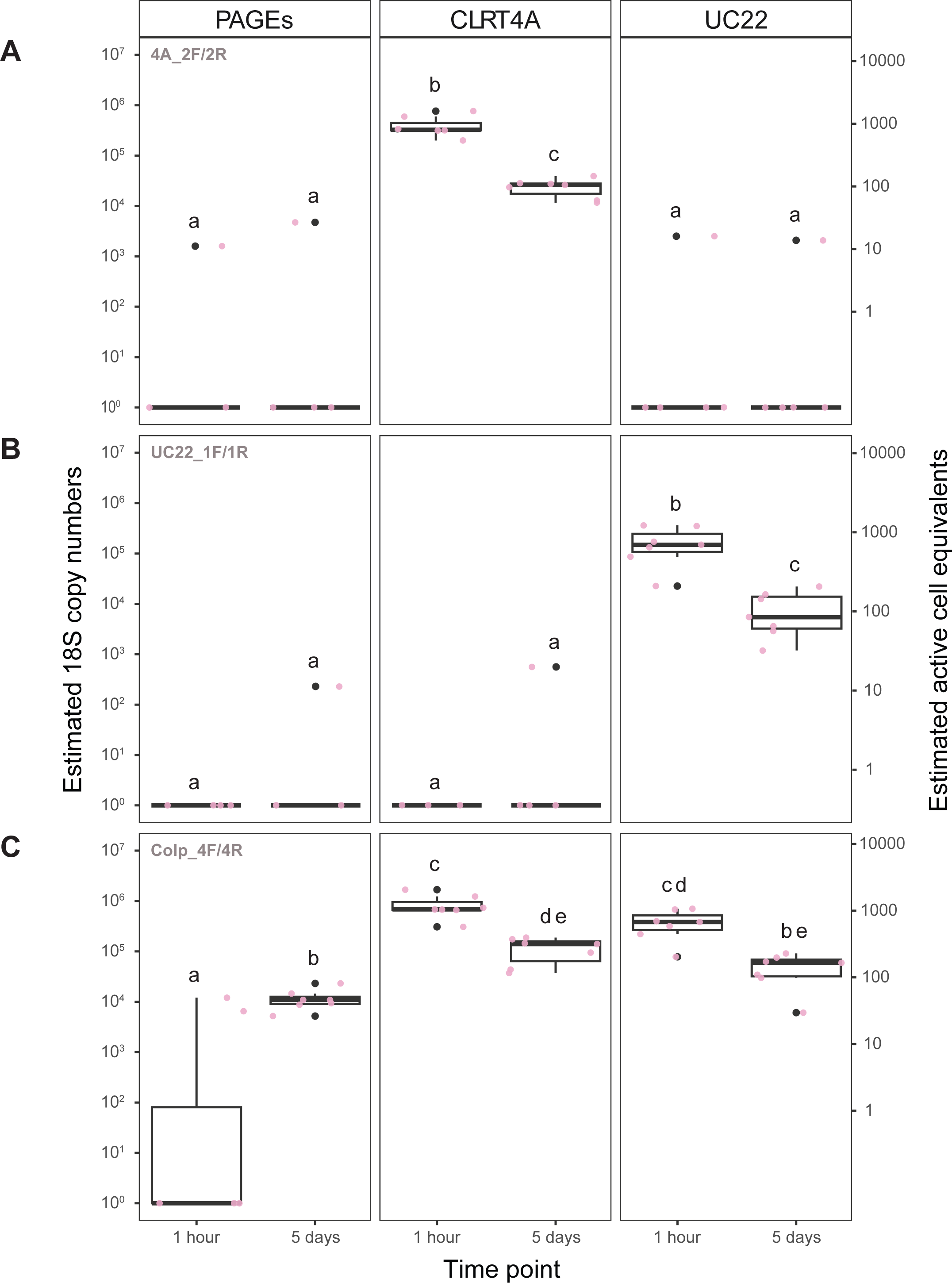
Boxplots showing the estimated quantities of Colpodida 18S fragments as well as estimated cell numbers in tomato plant leaflets inoculated with sterile PAGE’s solution, *P. steinii* (CLRT4A) from the phyllosphere, or *C. inflata* (UC22) from the rhizosphere. Letters above boxplots indicate statistical significance, as treatments with the same letters do not significantly differ. **A**. Fragment and cell numbers estimated when using the CLRT4A specific primers 4A_2F/4A_2R. **B**. Fragment and cell numbers estimated when using the UC22 specific primers UC22_1F/UC22_1R. **C**. Fragment and cell numbers estimated when using the broad Colpodida primers All_colp_4F/All_colp_4R.

Colpodida was undetectable on control plants when specific primers were used, and was generally low as determined by the broad range primers (Fig. 4C). The results indicate that the qPCR tests provide accurate estimates of the active inoculated protist populations on greenhouse plants, without significant interference from environmental organisms.

Estimated *P. steinii* and *C. inflata* populations declined by ∼90% in the five days after inoculation, with a mean ACE of ∼100 cells on each inoculated leaf for both organisms (p_adj_ < 0.001 with CLRT4A and UC22 specific primers, Fig. 4A-C, Table S5). *P*. *steinii* and *C. inflata* were detected on only one control leaf each at low abundances (< 100 cells). Low-level detection of Colpodida by the broad-specificity Colp_4F/4R primers increased on control plants over five days to a mean of ∼50 per leaf (p_adj_ < 0.001; Fig. 4C, Table S5). The nonspecific primers also detected significantly higher populations than the specific primers on *P. steinii-*inoculated plants (t_6_ = 4.533, p = 0.002; Fig. 4C). This could suggest that small populations of environmental Colpodida may have independently colonized the leaves. Together, these results demonstrate that Colpodida populations can be replicably modified on the leaf surface by inoculation, and that inoculated organisms are consistently detected after five days.

### Colpodida were ubiquitous on field-grown tomato leaves

To determine the range of absolute Colpodida abundance, we used qPCR to estimate Colpodida abundance on upper and lower leaves of tomato plants during dry and rainy conditions in the field. The phyllosphere clade of *P. steinii* was detected on all plants in wet conditions, including all but one upper leaf (Fig. 5A), but mostly limited to lower leaves in dry conditions. A 12-hour rainfall event significantly increased *P. steinii* population sizes, from a median 33 to 200 ACE/g fresh leaf weight on bottom leaves and from 0 to 166 on top leaves (Fig. 5A, values and statistics in Table S6, S7). In contrast, *C. inflata* was detected only on two or three lower leaves in either condition, and only once on an upper leaf (Fig. 5B, Table S6). This indicates that *C. inflata* does not exhibit the wetness-induced population increase seen in *P. steinii* on the leaves.

**Figure 5:**
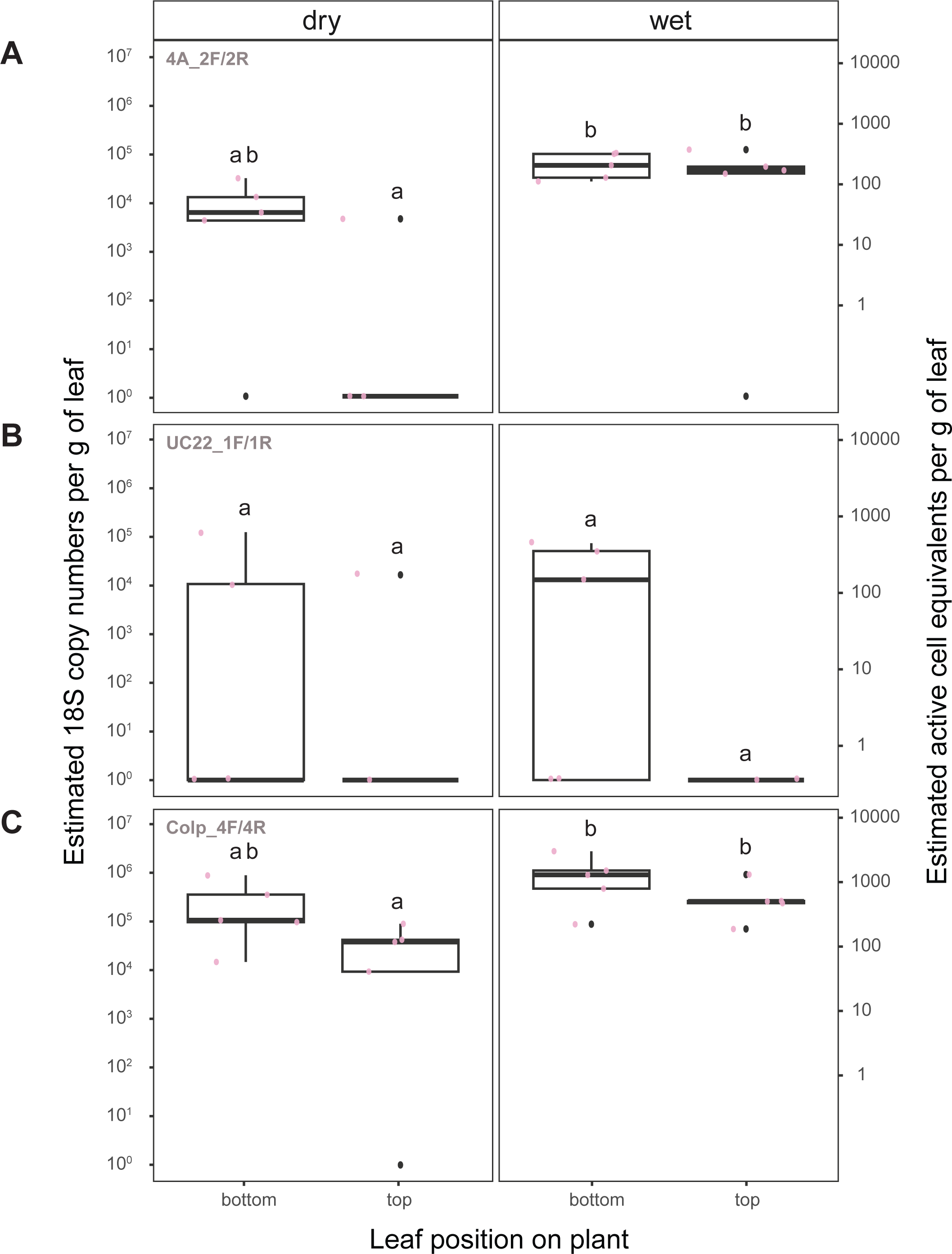
Boxplots showing the estimated quantities of Colpodida 18S fragments as well as estimated cell numbers on leaves of field-grown tomatoes collected on dry or wet days (i.e., after a rainfall). Leaves were collected from the tops or bottoms of plants. **A**. Fragment and cell numbers estimated when using the CLRT4A specific primers 4A_2F/4A_2R. **B**. Fragment and cell numbers estimated when using the UC22 specific primers UC22_1F/UC22_1R. **C**. Fragment and cell numbers estimated when using the broad Colpodida primers All_colp_4F/All_colp_4R.

Total Colpodida populations were detected using the Colp_4F/4R primer set in all but one leaf (Fig. 5C). Colpodida were detected on all plants, and similarly to *P. steinii*, populations increased significantly after rainfall and were greatest on lower leaves (Table S7). Median

Colpodida ranged from 122 ACE/g on top leaves in dry conditions to 1361 ACE/g on bottom leaves in wet conditions, significantly higher than estimated populations of *P. steinii* (t_19_ = 3.665, p = 0.002). This indicates that the phyllosphere *P. steinii* clade represents only one of multiple genotypes of Colpodida on tomato leaves. The *P. steinii* clade became more dominant under wet conditions, increasing from 8.4% to 18.7% of estimated Colpodida active cell equivalents after the rainfall event.

These results confirm that a clade of *P. steinii* protists comprise a significant minority of the ubiquitous Colpodida communities in the tomato phyllosphere, and show that their populations are highly responsive to wetting events and leaf position. *C. inflata* did not increase after rain, suggesting less adaptation to the phyllosphere niche. Together, the results validate the isolates including CLRT4A as representatives of common tomato phyllosphere protists in Connecticut, and demonstrate that Colpodida form prevalent, abundant, and moisture-responsive populations on tomato leaves.

## DISCUSSION

Heterophagous protists could play an influential role in structuring and activating the phyllosphere microbiota that are critical for plant immune development and resilience (Heil et al. 2025; Stone et al. 2018). Although previous work pointed to members of Colpodida as widely significant phyllosphere residents (Mueller and Mueller 1970, Taerum et al. 2023), an understanding of the identity and native abundance of these organisms was needed to confirm these hypotheses and enable future work. By studying newly developed isolates and PCR tools, this study confirms that Colpodida are consistently abundant in the phyllosphere of plants in our research plots, with dynamic populations affected by leaf age and wetness, and that a specific clade of *P. steinii* comprises up to one-fifth of the populations. The protocols and isolates presented here are available for future investigations of predator function, impact, and survival.

Although *P. steinii* was previously described from soil and permafrost environments (Li et al. 2024; Shatilovich et al. 2025), our results demonstrate that certain genotypes of *P. steinii* frequently colonize leaves. Isolation efforts favored a clade of *P. steinii* not represented by previous isolates; corresponding to a highly prevalent Colpodida ASV previously detected in leaf samples (Taerum et al. 2023). Members of this clade were also detected on almost all sampled leaves in the field after rainfall. In contrast, a rhizosphere-associated *C. inflata* clade was detected less frequently and mainly limited to lower leaves. Although both species have been described as “cosmopolitan” organisms able to inhabit diverse freshwater and soil environments, these observations suggest lifestyle specialization: the *P. steinii* clade has an ability to rapidly increase or activate its population on leaves after wetness, while *C. inflata* does not. This echoes the observations of Mueller and Mueller (1970), who observed that collected leaf water, but not rainwater, supported a rapid increase in a single dominant *Colpoda* morphotype. The authors proposed that nutrients and bacteria derived from the leaf were essential factors for protist growth. Our observation that *P*. *steinii* increases in dominance in wet conditions, and grows to higher yield than *C. inflata* in the presence of foliar pathogens, suggests that these leaf factors may also select some protists above others. Although we observed a decline in both *P. steinii* and *C. inflata* active cells after greenhouse inoculation, it could be that *P. steinii* has additional environmental requirements such as extended wetting events, certain bacterial prey, or abaxial locations for growth. More work is needed to develop inoculation methods, but based on field detection patterns and prior surveys, we hypothesize that the primary *P. steinii* clade isolated in this study represents a uniquely well-adapted leaf protist. The qPCR methods developed in this study can be used to test this hypothesis across different plant species and geographical locations.

The absolute abundance of predators on plant surfaces has rarely been investigated. We detected an estimated 10^2^-10^3^ Colpodida active cell equivalents of per gram leaf tissue in the field, which is on the same order of magnitude as estimates for leaf fungi (Leach et al 2017) and *C. cucullus* (Bamforth 1973). These numbers are dwarfed by population sizes of phyllosphere bacteria, which can reach over 10^6^ cells per gram (Williams and Marco 2014). However, Colpodida individuals in culture have been reported to consume between 10 and 200 bacterial cells per minute at high prey densities (Iriberri et al. 1995). Therefore, it seems likely that a measured population of 1000 active Colpodida individuals, with a theoretical feeding capacity of 10^4^-10^5^ prey per minute, could dramatically affect bacterial density, composition, and activity on a leaf, even if activity is limited to transient periods of wetness. Given the selective nature of predation and the ubiquity of Colpodida detection in the field, it is feasible that protist predation could be a source of selection pressure for bacteria in the phyllosphere as has been shown in soil bacteria (Nair et al. 2019), or similarly affect pathogen outcomes.

Many questions remain about the microeukaryotes making up the “microbial dark matter” of the phyllosphere. The isolates and quantification methods described in this study may serve as valuable tools to study *P. steinii* and other Colpodida across plant and environmental microbiomes, or adapted to study the population and community ecology of other plant-associated protists. A caveat is that the qPCR method is more effective at quantifying active organisms than cysts, and cannot distinguish encystment from true population decline. Counts may also be imprecise among active cells, as cells of *C. steinii* and *C. inflata* decrease in 18S copy numbers from peak activity to resting cyst stages (Zou et al. 2021), therefore quantitative estimates may depend on the life stage of the protists. Nevertheless, active populations have the closest relevance to protist function, and quantification will be informative in understanding in the colonization patterns, biogeography and host range, survival mechanisms, biological impact, and agricultural applications of Colpodida on plant surfaces.

## ACKNOWLEDGEMENTS

This research was funded by an AFRI Foundational Program grant from the United States Department of Agriculture National Institute of Food and Agriculture (USDA-NIFA 2022-67013-37144) to L.R.T., B.S., and S.J.T. We would like to thank Joe Liquori for assistance with growing and maintaining the tomato plants in the greenhouse experiment, Richard Cecarelli for growing the tomatoes on Lockwood Farm used in the field study, Quan Zeng for isolates of *Dickeya* and *Erwinia*, Johan Leveau for *Pantoea agglommerans* Pa299R, and Cole Wilson for technical laboratory assistance.

