## Supplementary figures and images for "A qPCR method facilitates study of absolute abundance, ecology, and inoculation fate of ciliate predators on the leaf surface"

### Fig. S1

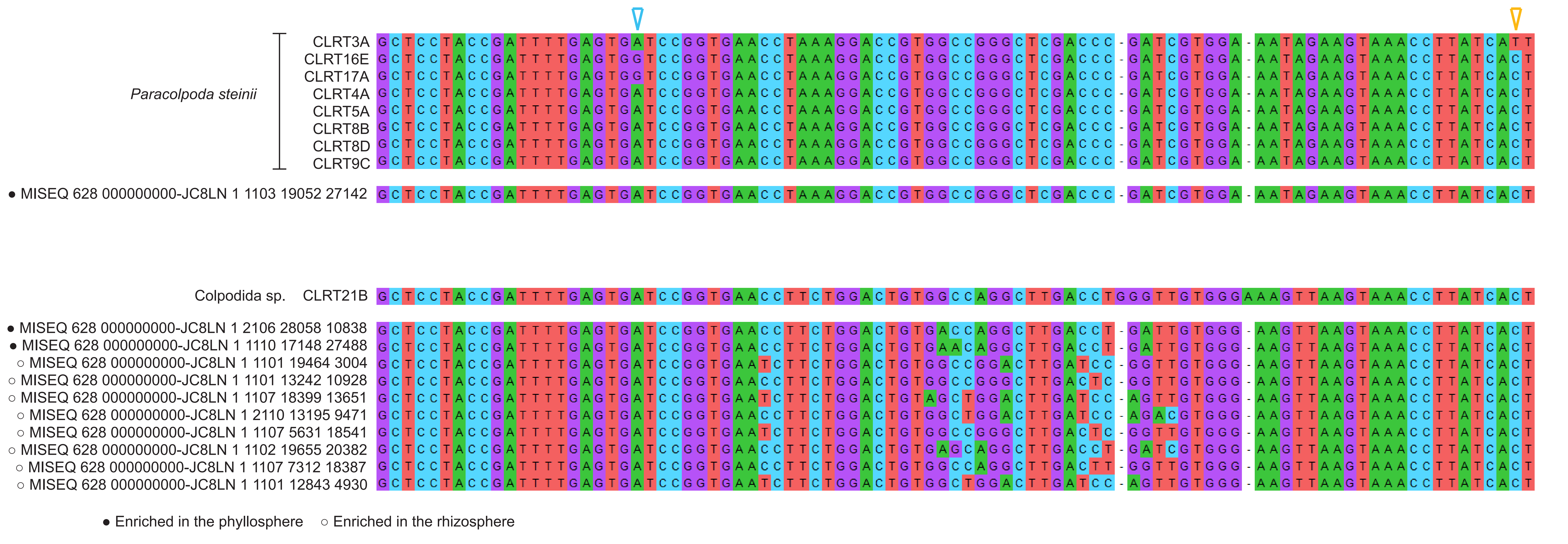

### Fig. S2

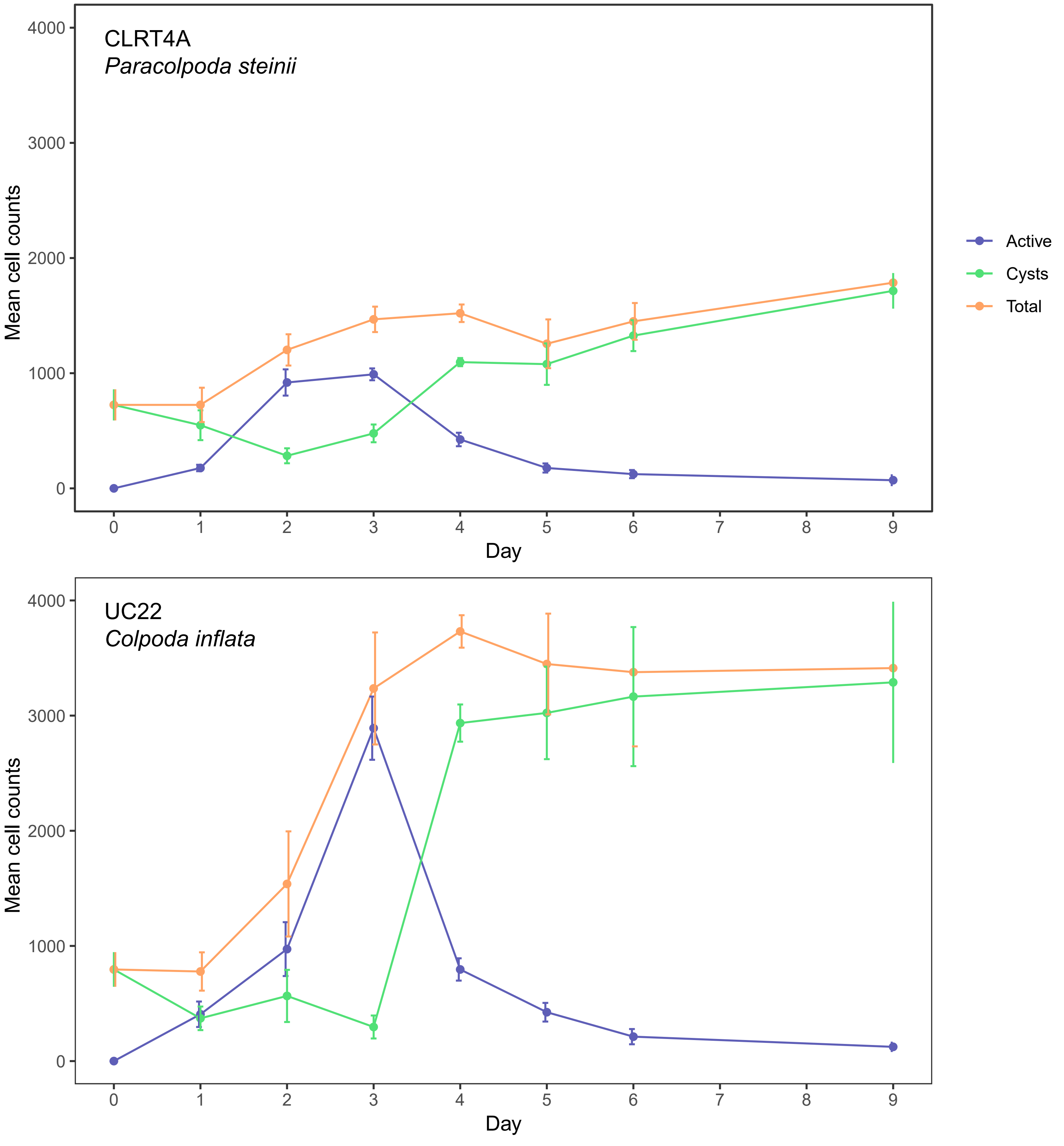

### Fig. S3

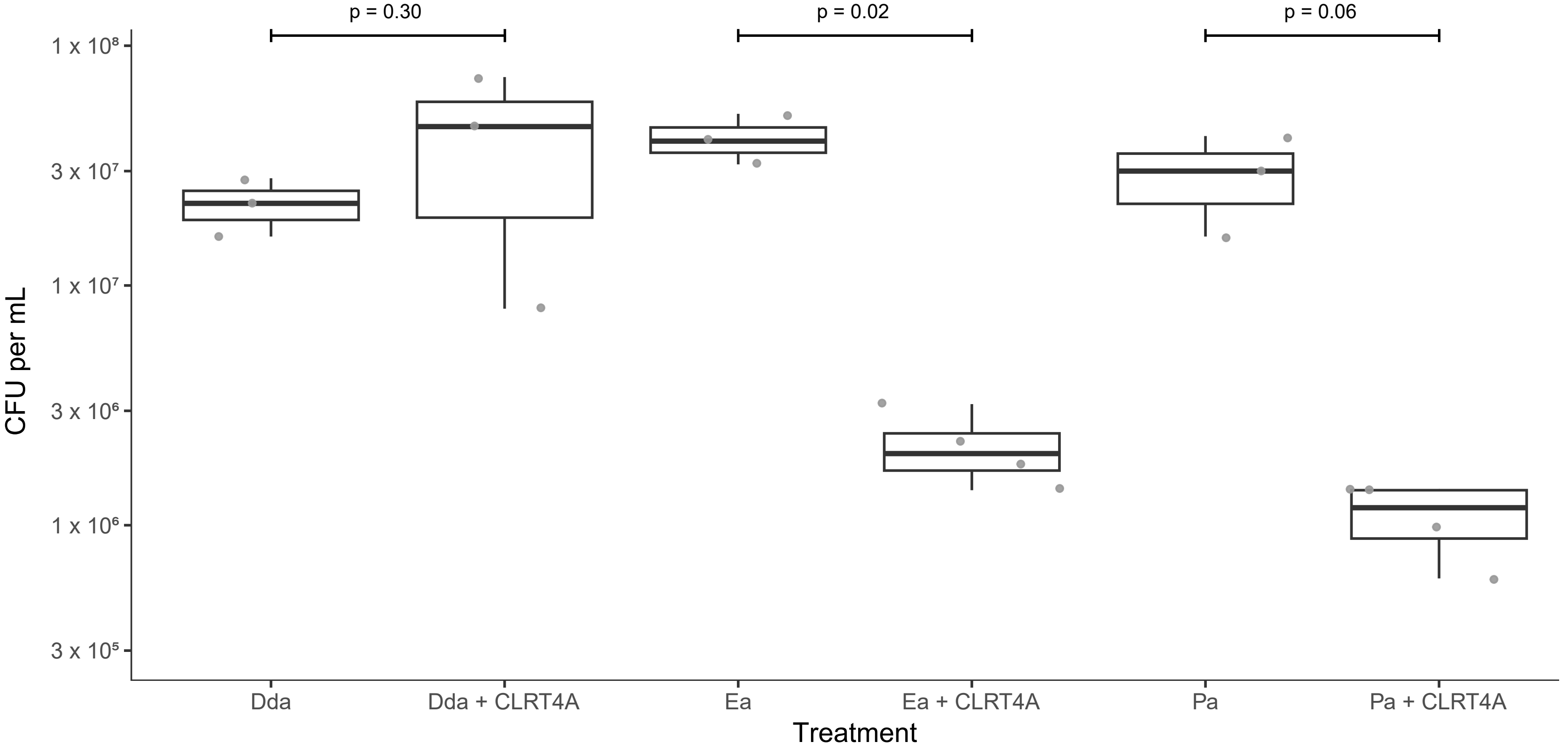

### Fig. S5

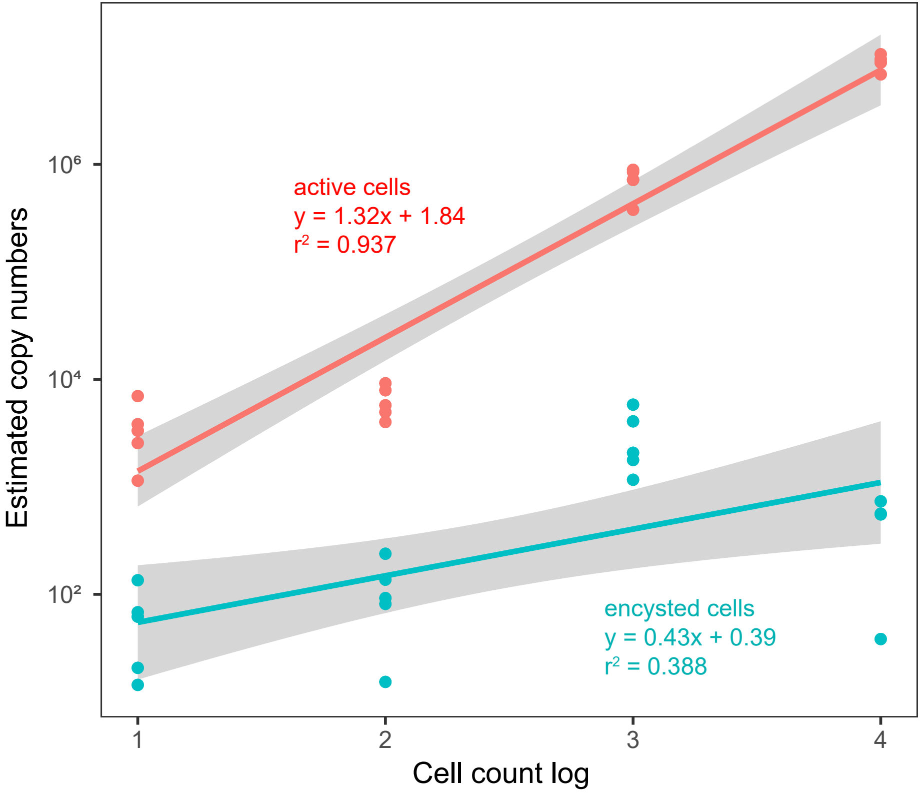
