## Supplementary material for "A qPCR method facilitates study of absolute abundance, ecology, and inoculation fate of ciliate predators on the leaf surface": Fig. S4

### Slide 1
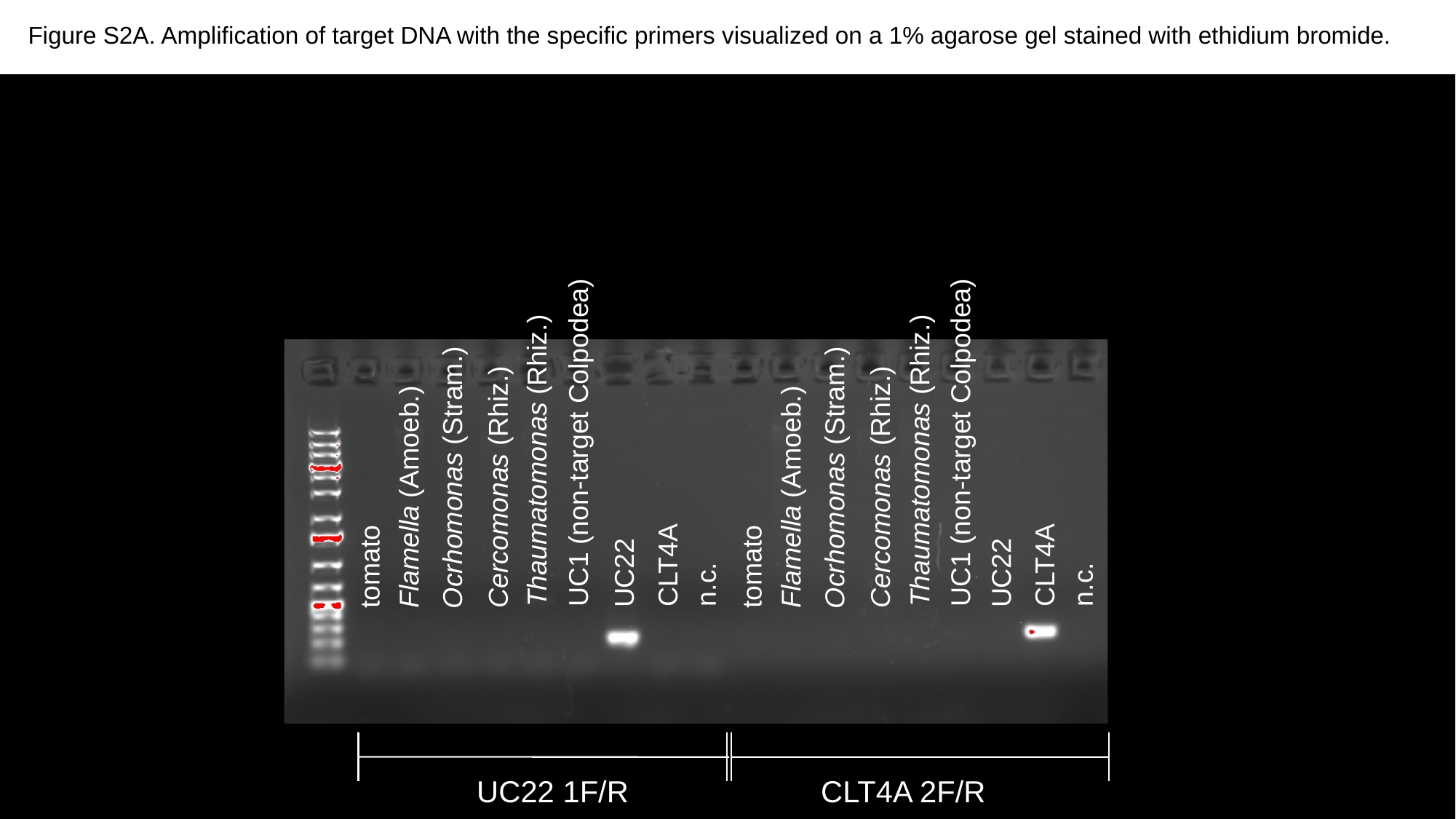

Figure S2A. Amplification of target DNA with the specific primers visualized on a 1% agarose gel stained with ethidium bromide.
UC1 (non-target Colpodea)
UC1 (non-target Colpodea)
Thaumatomonas (Rhiz.)
Thaumatomonas (Rhiz.)
Ocrhomonas (Stram.)
Ocrhomonas (Stram.)
Cercomonas (Rhiz.)
Cercomonas (Rhiz.)
Flamella (Amoeb.)
Flamella (Amoeb.)
CLT4A
CLT4A
tomato
tomato
UC22
UC22
n.c.
n.c.
UC22 1F/R
CLT4A 2F/R

### Slide 2
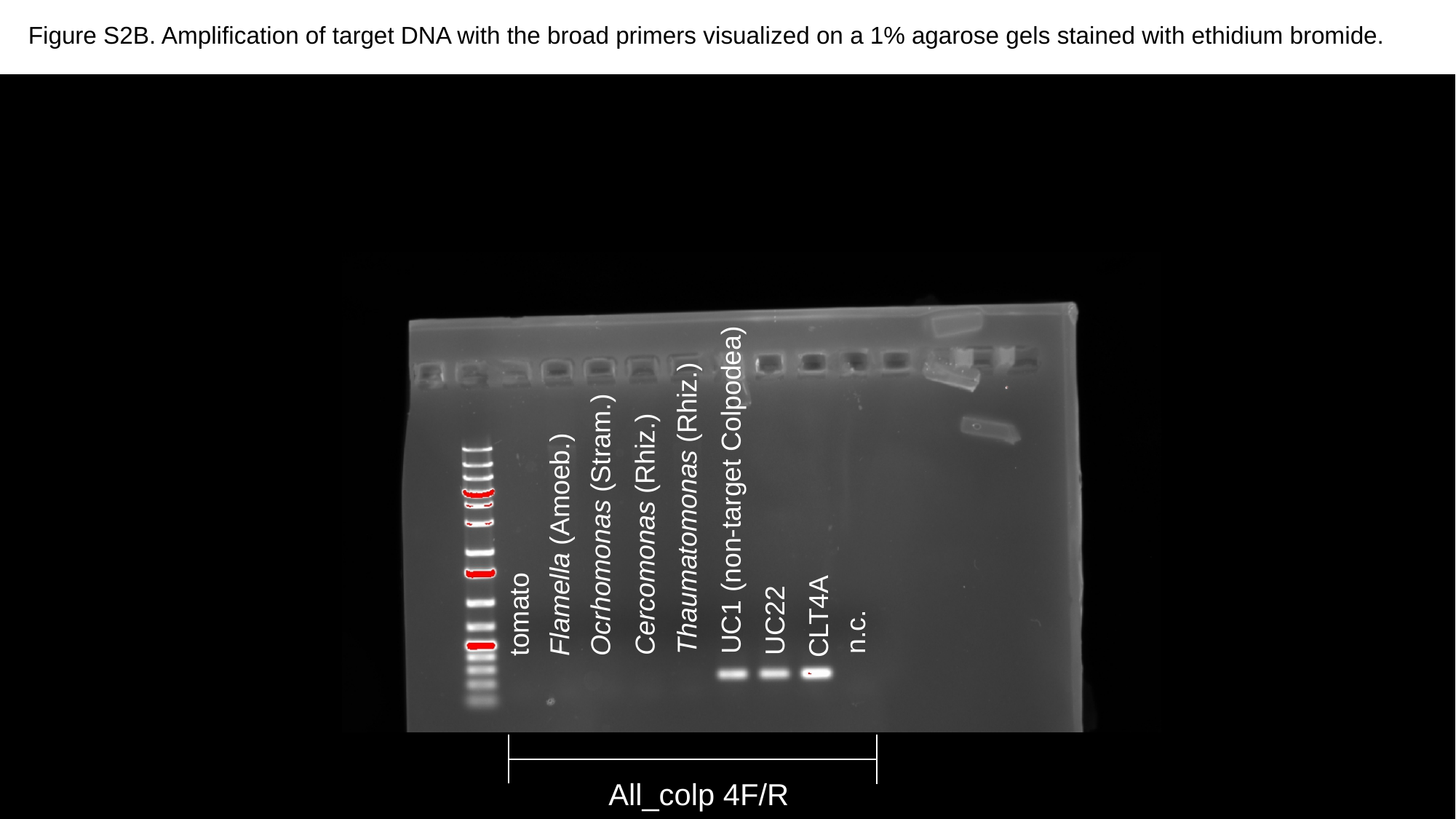

Figure S2B. Amplification of target DNA with the broad primers visualized on a 1% agarose gels stained with ethidium bromide.
UC1 (non-target Colpodea)
Thaumatomonas (Rhiz.)
Ocrhomonas (Stram.)
Cercomonas (Rhiz.)
Flamella (Amoeb.)
tomato
CLT4A
UC22
n.c.
All_colp 4F/R
